# MetaMLP: A fast word embedding based classifier to profile target gene databases in metagenomic samples

**DOI:** 10.1101/569970

**Authors:** G. A. Arango-Argoty, L. S. Heath, A. Pruden, P. J. Vikesland, L. Zhang

## Abstract

The functional profile of metagenomic samples allows the understanding of the role of the microbes in their environment. Such analysis consists of assigning short sequencing reads to a particular functional category. Normally, manually curated databases are used for functional assignment where genes are arranged into different classes. Sequence alignment has been widely used to profile metagenomic samples against curated databases. However, this method is time consuming and requires high computing resources. Although several alignment free methods based on k-mer composition have been developed in the recent years, they still require a large amount of memory. In this paper, MetaMLP (Metagenomics Machine Learning Profiler) a machine learning method that represents sequences into numerical vectors (embeddings) and uses a simple one hidden layer neural network to profile functional categories is proposed. Unlike other methods, MetaMLP enables partial matching by using a reduced alphabet to build sequence embeddings from full and partial kmers. MetaMLP is able to identify a slightly larger number of reads compared to Diamond (one of the fastest sequence alignment method) as well as to perform accurate predictions with 0.99 precision and 0.99 recall. MetaMLP can process 100M reads in around 10 minutes in a laptop computer which is 50x faster than Diamond. MetaMLP is free for use, available at https://bitbucket.org/gaarangoa/metamlp/src/master/.

## Introduction

The wide and rapid adoption of next generation sequencing techniques (NGS) such as metagenomics in the analysis of microbial diversity, antibiotic resistance, and other functional profiling analysis creates a gap between scalability and processing efficiency. In other words, large amounts of data require the design of computational tools that are both accurate and fast. Sequence comparison algorithms such as BLAST (1), FASTA (2), HMMER (3), PSI-BLAST (4) were created with the aim to find correspondence of the sequence distribution in two or more sequences. BLAST is to date the most popular and trusted tool for sequence alignment. However, it is well known that BLAST does not escalate well when comparing millions of sequences. The reason is that BLAST uses a computationally demanding strategy consisting of a seed and extend algorithm (5). Although, sequence alignment is considered the gold standard approach for sequence analysis, there are several cases where this technique can produce dubious results (6). For instance, alignment-based methods assume that homologous sequences share a certain degree of conservation. Although this assumption is considered to be true when analyzing conserved domains, organisms such as viruses that exhibit a high degree of mutation, challenge this collinearity principle. When analyzing short sequences (e.g., Illumina sequencing reads), the percentage of identity does not guarantee correctness. Highly identical sequences do not imply homology (7). In the opposite case, sequences with less than 30% identity can potentially be considered as homologous (8).

DIAMOND (9), BLAT (10), USEARCH (11), and RAPSearch (12) are alternatives to BLASTX that can run much faster but with a loss of sensitivity. Particularly, the dramatic speed up of DIAMOND its (20,000X) is achieved by using a double indexing strategy, spaced seeds (longer seeds where not all positions are used) and a reduced alphabet. In detail, DIAMOND implements a seed and extend algorithm that first indexes both query and reference sequences. Then, the list of seeds in both the query and reference are linearly traversed to determine all the matched seeds with their locations. Finally, seeds are extended by using the Smith-Waterman algorithm (13).

Alignment-free methods have been proposed as an alternative to quantify the sequence similarity without performing any sequence alignment (6,14). These methods do not use the seed and extend paradigm. Therefore, their computational complexity is often linear and only depends of the query sequence length. In next-generation sequencing, several alignment-free strategies have been developed for different applications, including transcript quantification (kallisto (15), sailfish (16), Salmon (17), RNA-Skim (18)), variant calling (ChimeRScope (19), FastGT (20)), de-novo genome assembly (minimap (21), MHAP (22)), and the profiling of metagenomics taxonomy by using a kmer matching approach (Kraken (23), Mash (24), CLARK (25), stringMLST (26)).

The word embeddings technique is one of the most successful learning methods applied in natural language processing (NLP) where words can be represented as a numerical vectors. For instance, the Word2vec technique (27) uses a shallow two-layer neural network to train and aggregate word embeddings by using the continuous bag of words (CBOW) approach. Thus, identifying semantic associations between a target word given its context. The concept of using word vectors for representing protein/DNA sequences is not new and has been explored before. For instance, DNA2Vec (28), explores the associations between variable length kmers to generate an embedding space that proved to correlate with sequence alignment. Yang et. al., (29) explores the performance of word embeddings for classification of protein functions compared with classical representation techniques. Yan et. al., demonstrated that kmer embeddings outperformed other techniques. However, in both studies, embeddings are learnt in an unsupervised way. This means, that the embeddings are learnt first and then the classifier is built by using those embeddings. In this paper, MetaMLP, an alignment-free method that uses word embeddings to represent target protein databases is proposed for the functional profiling of metagenomic samples. The strategy behind MetaMLP relies on the CBOW model. However, the target word is replaced by the label or functional class of the sequence and the context words corresponds to the kmers and fragmented kmers. Therefore, MetaMLP is a novel strategy that uses a combination of hash indexing, six open reading frame translation, a reduced amino acid alphabet and an embedding representation to process metagenomic data. In addition, MetaMLP was built up on top of the C++ FastText (30) library and its composed of two main stages: MetaMLP-index that process protein sequences to build a machine learning model and MetaMLP-classify to annotate reads from metagenomic DNA sequence libraries.

## Methods

The overall structure of MetaMLP is shown in Figure 1 and consists of two main components: A) An indexing stage that process protein reference sequences into a word vector representation to train a classifier and B) A prediction stage that process short sequencing reads and classifies them into one to the predefined classes from the reference database.

**Figure 1:**
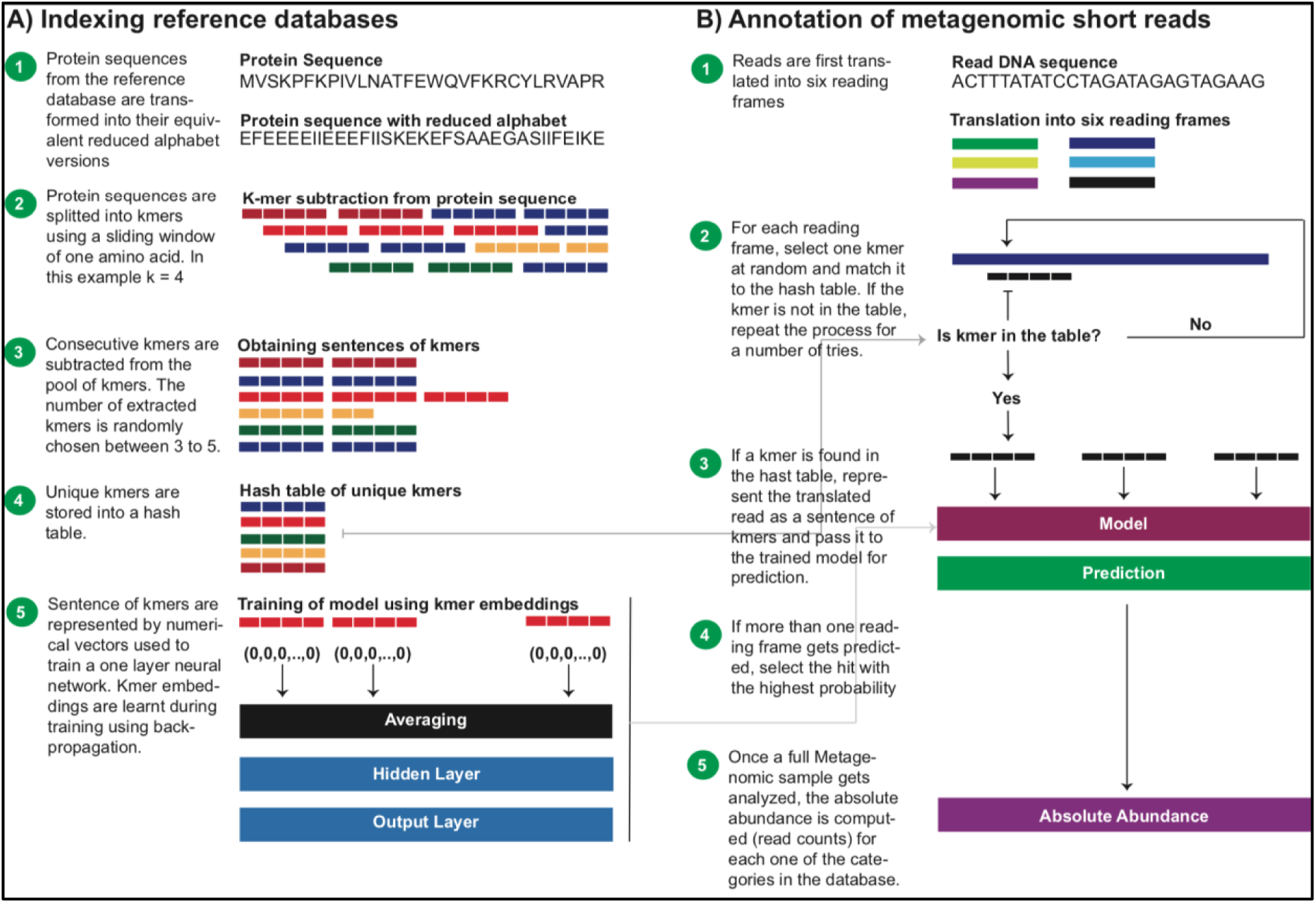
Overview of MetaMLP. A) Indexing reference databases where proteins are used to train the machine learning model. B) Once a model is trained, it will be used later to profile short sequencing reads to produce a relative abundance profile and the individual predictions for each read.

### Indexing Protein Reference Databases

#### Reference Database Preprocessing

To increase the chances of detecting sequences with mismatches, the reference proteins are first transformed into their equivalent 10 amino acid alphabet version using the murphy.10 alphabet representation used in rapsearch (A [KR] [EDNQ] C G H [ILVM] [FYW] P [ST]) (31). Then, kmers of a fixed length are extracted from each protein sequence. However, to consider all kmers within a sequence, a sliding window of one amino acid is used. Thus, each protein comprises k versions, each one corresponding to a different starting location [1,…, k]. Thereafter, a ‘sentence’ of kmers is extracted by taking 3 to 5 consecutive kmers (equivalent of reads of 100 to 150 bp) (see Figure 1A). At the same time, a table with unique kmers is built and stored to later be used for filtering sequences that diverge greatly from the reference database during the prediction stage.

#### Training

MetaMLP uses the FastText implementation of the continuous bag of words (CBoW) technique to learn the semantic relations between protein sequences and their labels. Thus, proteins are represented as a series of kmer sentences (analog to sentence of words in text documents). Then, it decomposes each kmer within the sentences into a numerical representation (kmer vector) (see Figure 1A). Later, it computes the average of the kmer vectors and passes it to a single hidden layer neural network. Finally, it outputs the probability distribution over the established classes by using a softmax layer. In addition, MetaMLP enables the bag of n-grams feature from FastText to capture partial information from the kmers. These n-grams are sub sequences from the kmers passed along with the full size kmer allowing to identify kmers with partial matching.

#### Prediction of Short Sequencing Read

MetaMLP is designed to efficiently profile metagenomic samples with millions of reads from short sequencing libraries against a target reference database. As reads are made of nucleotides, MetaMLP first translates each sequence into six reading frames. Then, for each reading frame a random kmer is selected from its sequence and checked against the hash table that was built during the indexing stage. If a kmer is found in the hash table, all kmers are subtracted from the read and classified using the trained CBoW model. If not, a new random kmer is selected from the read at a different position. This process is repeated to a maximum number of tries defined by the user. If more than one reading frame gets classified, MetaMLP picks up the reading frame with the highest classification probability (see Figure 1B).

Once a full metagenomic dataset is processed, MetaMLP counts the number of reads per class using a minimum probability cutoff defined by the user and reports the absolute abundance table. Additionally, MetaMLP also reports a fasta file containing the read name along with its classifications, probabilities and sequence. This file is useful for cases where MetaMLP is used as a filter to target a particular functional classes.

### Databases

#### Pathway Reference Database

Bacterial protein sequences from the Universal Protein Resource (UniProt) were downloaded and filtered by only proteins that have been manually curated, reviewed and contained evidence at the protein level. In total 20,161 proteins were obtained and 4,105 of those were annotated to at least one pathway. Lastly, pathways with less than 50 proteins were discarded to get a total of 3,216 proteins and 21 different pathways (see Supplementary Table S1).

#### Antibiotic Resistance Database

MetaMLP was trained to identify short reads associated to Antibiotic Resistance Genes (ARGs) from metagenomic short sequencing data. Thus, the DeepARG-DB-v2 database (32) containing a total of 12,260 sequences distributed through 30 antibiotic categories was downloaded. However, only antibiotic resistance categories with at least 50 protein sequences were considered for downstream analysis. Thus, a total of 12,147 proteins and 14 categories were used to train the MetaMLP model (see Supplementary Table S2).

#### Gene Ontology Reference Database

Protein sequences associated to the biological process response to stress (GO:0006950) were downloaded from UniProt website. However, only bacterial curated sequences and biological processes with at least 100 sequences were considered for downstream analysis (see Supplementary Table S3). In addition, the GO database comprises proteins with multiple associated labels. For instance, the protein sequence Q55002 is associated to response to antibiotic (GO:0046677) and translation (GO:0006412). Therefore, reads from this protein would be classified to both categories. However, as MetaMLP uses a softmax layer for prediction, it will distribute the probability between both categories. In an ideal scenario, both classes would have a probability of 0.5. This database was used to test the ability of MetaMLP to represent sequences associated to multiple labels.

### True Positive Dataset

The pathway database was used to build a true positive database. Because MetaMLP uses amino acid sequences for training and nucleotide sequences for querying, it was necessary to identify the corresponding nucleotide sequences for each one of the proteins in the pathways database. Therefore, UniProt identifiers were cross referenced against the RefSeq database and a list of gene candidates were found. Then, those candidates were aligned to the protein sequences using diamond BlastX with a 90% identity and a 90% overlap. If multiple alignments were obtained at this criteria, the best hit was selected as the representative gene sequence for the target protein sequence. Thus, each entry in the database contained a respective gene sequence. Finally, the pathway database was randomly splitted into training (80%) and validation (20%). The training set was used to tune the model whereas the validation set was exclusively used to test the method after the training was done. Therefore, the validation set was never used during training process. Note that the training set corresponds to amino acid sequences whereas the validation set consists of nucleotide sequences. To simulate a library of short sequence reads, sequences of 100bp long were randomly subtracted from each nucleotide sequence from the validation dataset. Thus, a total of 35,751 short reads were generated.

Diamond is currently one of the widely used tools for metagenomic analysis. Therefore, to test the performance of MetaMLP, diamond BlastX with the best hit approach was used. Diamond was run by using a sequence alignment identity of 80%, whereas MetaMLP was set with a minimum probability of 0.8. Precision, Recall and F1 score were computed to measure the performance of both approaches.

### False Positives Dataset

To test the ability of MetaMLP to filter out sequences that are not associated to any of the selected pathways (false positives), a synthetic dataset was constructed by using the same number of reads from the true positive dataset. However, each nucleotide position on this dataset was randomly selected. This negative dataset was then ran against MetaMLP and the best hit approach using Diamond with default parameters. Precision, recall and F1 score were computed to measure the performance of both methods.

### Time and Memory Profiling

To evaluate the time performance and memory footprint of MetaMLP, a dataset of 100k, 1M, 10M and 100M reads were built by randomly subtracting reads from a real metagenomic soil sample of 407,645,066 reads. This sample is under the SRA accession number SRR2901746 and corresponds to a 250bp long read sample from the Illumina HiSeq 2000 sequencer. Along with MetaMLP, diamond was also ran with the same datasets. Both methods ran with only one enabled CPU in the same linux 16.4 environment.

### Functional annotation of Metagenomic Datasets

MetaMLP was used to profile four different environments comprising a total of 68 metagenomic samples through the functional composition analysis including: Pathways detection, response to stress, and antibiotic resistance composition. The 68 public available metagenomes were downloaded from the Sequence Read Archive (SRA) from the National Center for Biotechnology Information (NCBI) spanning four different environments as follows: 10 soil (S), 15 human gut (HG), 15 freshwater (FW) and 28 wastewater (WW) samples (see Supplementary Table S4 for details). Results from MetaMLP were compared against the best hit approach using Diamond BlastX with an identity of 80%.

For the GO reference database, MetaMLP was run with a permissive 0.5 minimum probability to retrieve multiple classifications. Relative abundance results were compared against those obtained using sequence alignment with diamond BlastX at an 80% identity cutoff.

### Availability of MetaMLP

Source code for MetaMLP is available at https://bitbucket.org/gaarangoa/metamlp/src/master/

## Results and Discussion

The sequence embedding strategy allows MetaMLP to represent amino acid sequences into numerical vectors (embedding dimension) by taking into account the distribution of the kmers in the protein sequence as well as their labels. Thus, MetaMLP uses the supervised embedding implementation from FastText to learn these numerical vectors and minimize the inner distance within members of a class and maximize the outer distance to other classes. For instance, proteins that belong to Beta-lactamase class are expected to cluster together and keep distant from members of other classes. Figure 2 shows the distribution of the MetaMLP embeddings in a two dimensional space generated by using the t-SNE algorithm (33). For targeted databases such as the ARG categories or pathways database, MetaMLP clustered categories according to their labels with a representative cohesion and separation (silhouette score: 0.56 and 0.62 for pathways and ARGs, respectively) (See Figure 2A-B). Interestingly, in a complex classification problem represented by the GO database where proteins contain multiple labels, MetaMLP show a consistent distribution over the clusters and its corresponding categories. Clusters shown in Figure 2C describes the relationship among different biological processes involved in response to stress. For example, proteins responding to antibiotics are also associated to other biological process such as response to toxic substances, pathogenesis, defense to virus, chemotaxis, response to DNA damage, among others. Such associations can be clearly seen from the embeddings visualization. Therefore, the embedding strategy adopted in MetaMLP is also suitable for representing reference databases where proteins contains multiple labels.

**Figure 2:**
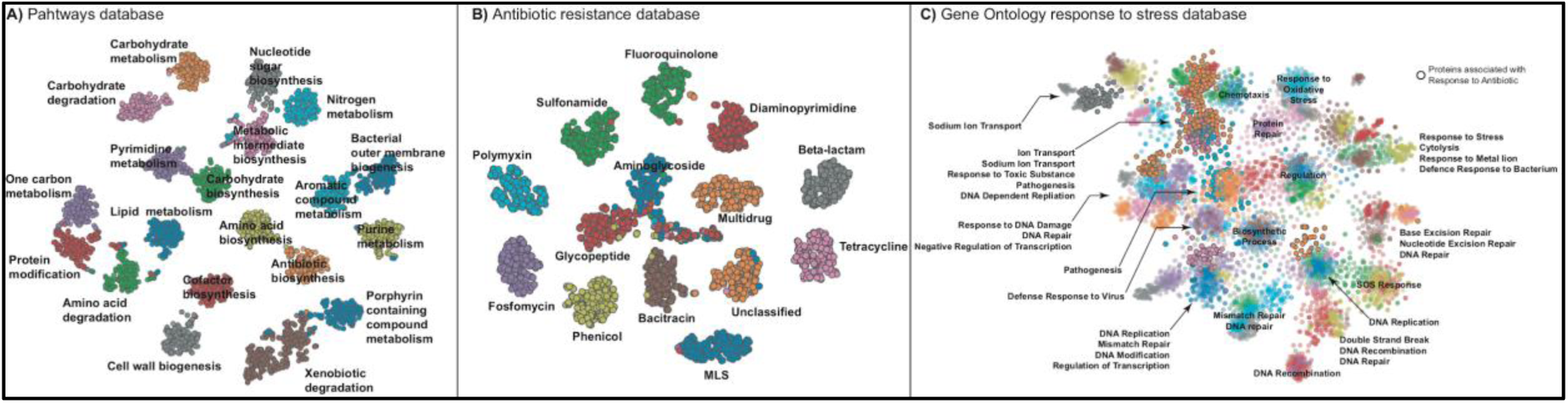
MetaMLP embeddings representation in two dimensional space for the pathways, ARGs and GO response to stress databases.

### Detection of True Positive Hits

The pathways database was used to assess the ability of MetaMLP to 1) discriminate between pathway-like reads and 2) to evaluate the performance of MetaMLP on classifying short sequences from a particular pathway. To compare the performance of MetaMLP, the best hit approach using Diamond BlastX was used. In total, MetaMLP was able to identify 10,433 (29%) pathway-like reads out of the total 35,751 with a probability greater than 0.8, whereas, the baseline approach was able to identify 8,695 (24%) reads out of the 35,751. This means MetaMLP was able to identify 5% more reads than the best hit approach at 80% identity. Further, both methods were compared on their positive predictions to evaluate their performance for discriminating reads from a particular pathway. As expected, the sequence alignment approach at 80% identity performed with a high average precision (0.99) and recall (1.00) (see Supplementary Table S5) whereas MetaMLP was also near to a perfect prediction with a 0.99 average precision and 0.99 average recall (see Supplementary Table S6) indicating the potential of the kmer vectors to represent protein sequences to profile metagenomes. It is also worth mentioning that MetaMLP and the best hit approach didn’t perform well for three categories (Aromatic compound metabolism, Bacterial outer membrane biogenesis, and xenobiotic degradation). Interestingly, the best hit approach was not able to identify any read from the bacterial outer membrane biogenesis when MetaMLP obtained a 1.00 precision but a low 0.13 recall indicating a high sensitivity of MetaMLP in discriminating true positives from this category but failing for false negatives. In terms of relative abundance, the comparison of the read counts between the best hit approach and MetaMLP was very close with a correlation of 0.988, indicating that MetaMLP can correctly characterize the composition of the pathways in the simulated dataset (see Supplementary Figure S1).

### Detection of False Positives Hits

A false positive is a negative sample predicted as positive. For instance, a read that does not belong to any pathway class is predicted to particular pathway. In this false positive scenario, MetaMLP was tested against the number of predicted random reads by counting how many out of the 35,751 negative reads were classified in any pathways. As result, MetaMLP classified only 2 reads (0.005%) out of the 35,751 negative reads indicating a very low false positive rate. As expected, the best hit approach didn’t produce any relevant alignment.

### Time and Memory Usage of MetaMLP

The main advantage for building up a classifier instead of performing a sequence alignment is the improvement over the speed for making the annotations. Results have shown that MetaMLP keeps an almost identical level of sensitivity compared to Diamond BlastX. However, the strength of MetaMLP relies on its speed. Table 1 shows the speed benchmarking over datasets with different number of reads. Note that MetaMLP is >50x times faster than Diamond for all the sample sizes. MetaMLP produces very similar results in terms of relative abundance using the ARGs database and pathway database with a correlation of 0.951 and 0.953, respectively (See Supplementary Figure S2). Note that in this test, MetaMLP identified 35% more ARG-like reads (253,370) compared to the number of reads (186,736) detected from Diamond BlastX. In addition, MetaMLP is also memory efficient depending mostly on the size of the reference database. For instance, it requires a minimum ram memory of 1.0Gb to run the pathways database, 1.2Gb when using the ARGs database and 2.8Gb if using the GO database. When processing 100M reads, it required 1.7Gb in total with the pathways database whereas Diamond BlastX required 6.68Gb. The low memory usage in MetaMLP is a consequence of its classification strategy where reads are loaded in chunks of 10,000 reads for efficient I/O rate. Therefore, MetaMLP can be run in any personal computer without the need of using a big cluster with high amount of RAM memory.

**Table 1:** Time profiling of MetaMLP compared to Diamond BlastX over different sample sizes.

| Number of Reads | MetaMLP | Diamond |
| --- | --- | --- |
| 100,000 | 9 s | 38 s |
| 1,000,000 | 27 s | 6 m |
| 10,000,000 | 1 m | 67 m |
| 100,000,000 | 14 m | 714 m |

### Functional Annotation of Different Environments

MetaMLP was run over the 67 real metagenomic samples processing a total of 2,186,933,071 reads. Of those reads MetaMLP was able to predict 2,343,026 as ARG-like reads in 710 minutes using only once CPU, whereas Diamond BlastX identified 2,003,050 reads taking a total of 5,256 minutes using 20 CPUs. Thereafter, the average correlation of the abundances between Diamond and MetaMLP was of 0.94 (0.88 log transformed abundance). Interestingly, human gut microbiota and wastewater were the two environments where both methods had the highest correlation respect their log transformed abundance (0.96, 0.93 respectively) whereas soil and freshwater had each a correlation of 0.83.

#### Observation of MetaMLP Annotations against an extensive Metagenomics Study

An extensive study carried out by Pal et. al., (34), uses over >800 metagenomic samples spanning several environments with a sequence alignment strategy at a 90% identity cutoff for annotation. This study (named Pal800 for simplicity) shown that the human gut microbiota is one of the environments with the highest relative abundance compared to other microbiomes (soil, wastewater and freshwater). Concordantly, when MetaMLP was run over the 68 real metagenomic samples using the GO database, it also profiled the human gut microbiome as the highest relative abundance for the response to antibiotic process (see Supplementary Figure S3). Note that Pal800 used a curated ARG database and therefore it didn’t consider the induction of false positives. However, the GO database only provides a general overview of the functional composition of those environments. Therefore, a more detailed analysis was obtained by looking at the results from MetaMLP using the specialized ARGs database. As result, the same trend was observed when comparing both analysis (MetaMLP, Pal800). For example, the tetracycline category had the highest relative abundance in the human microbiome, sulfonamide shows the highest relative abundance in the wastewater environment, the relative abundance of the beta-lactamase class was higher in the freshwater compared to the wastewater and both are higher than human gut and soil environments. Pal800 also performed a composition profile of the mobile genetic elements present in the microbiomes. It shown that wastewater, freshwater and soil environments had a higher relative abundance compared to the human gut. Interestingly, for MetaMLP the GO response to stress database conveyed a similar trend in relative abundance for the biological process “establishment of competence for transformation” (Transformation Supplementary Figure S3). This term is associated to genetic transfer between organisms and is described by the GO consortium as the process where exogenous DNA is acquired by a bacterium. In overall, despite only using 67 real metagenomes, the functional annotation carried out by MetaMLP described a very similar trending for relative abundances when compared to the Pal800 study indicating a real scenario usage of MetaMLP.

## Conclusions

MetaMLP is an alignment-free method for profiling metagenomic samples to specific target group of proteins (e.g., ARGs, pathways, GO terms) using a machine learning classifier. It uses sequence embeddings to represent protein/DNA sequences as numerical vectors and a linear classifier to discriminate between protein functions. Results show that MetaMLP identifies more reads than the widely used best hit approach (sequence alignment with identity >80%) and has a good performance as the sequence alignment method. Remarkably, MetaMLP is around 50x faster than DIAMOND aligner, the most widely used sequence alignment tool for metagenomic datasets. MetaMLP can be trained using any collection of protein sequences (reference database) and keeps a very low memory footprint for the specialized databases used in this paper. Finally, MetaMLP is open sourced and freely available at https://bitbucket.org/gaarangoa/metamlp/src/master/.

