## Supplementary material for "MetaMLP: A fast word embedding based classifier to profile target gene databases in metagenomic samples"

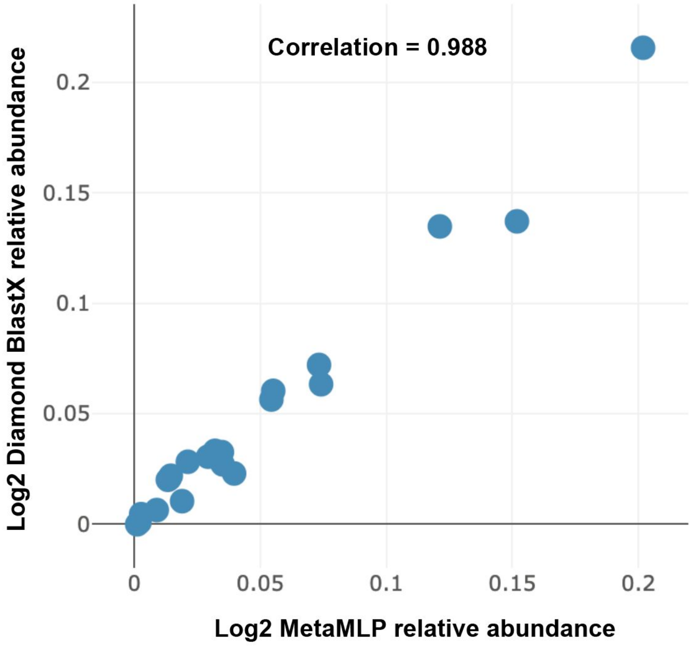

**Supplementary Figure S1**: Correlation between pathway relative abundances computed from results from MetaMLP (x axis) and Diamond BlastX (y axis) using the true positive dataset.

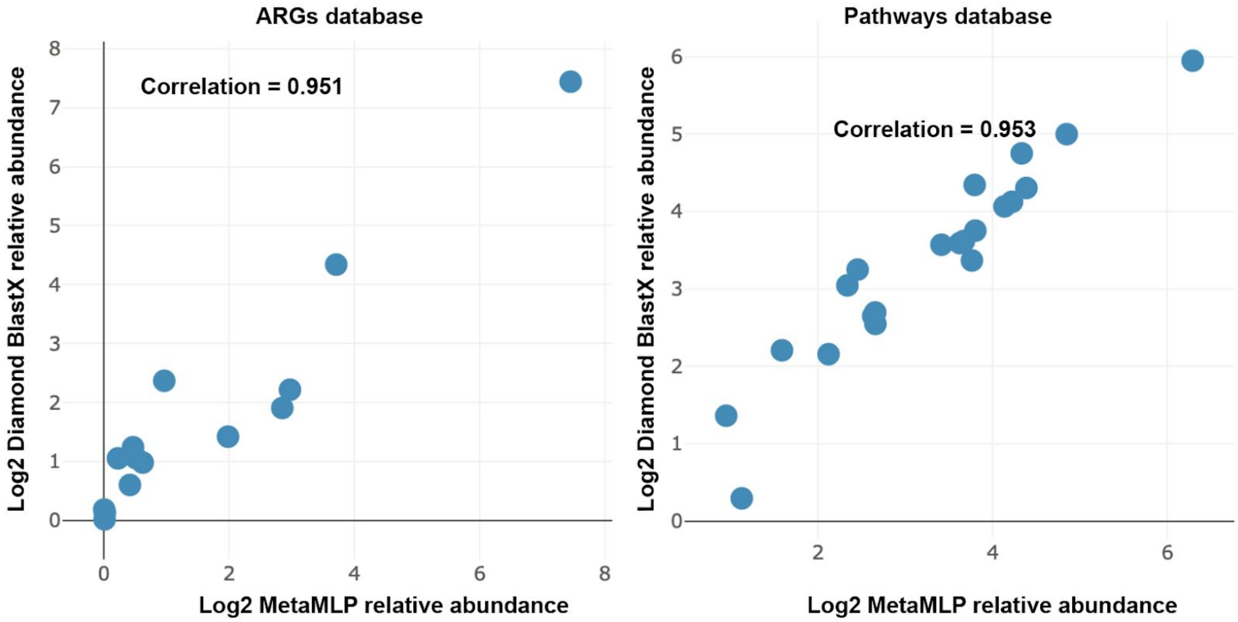

**Supplementary Figure S2**: Correlation between MetaMLP and Diamond BlastX relative abundance results from the 100 Million dataset obtained from a real soil sample.

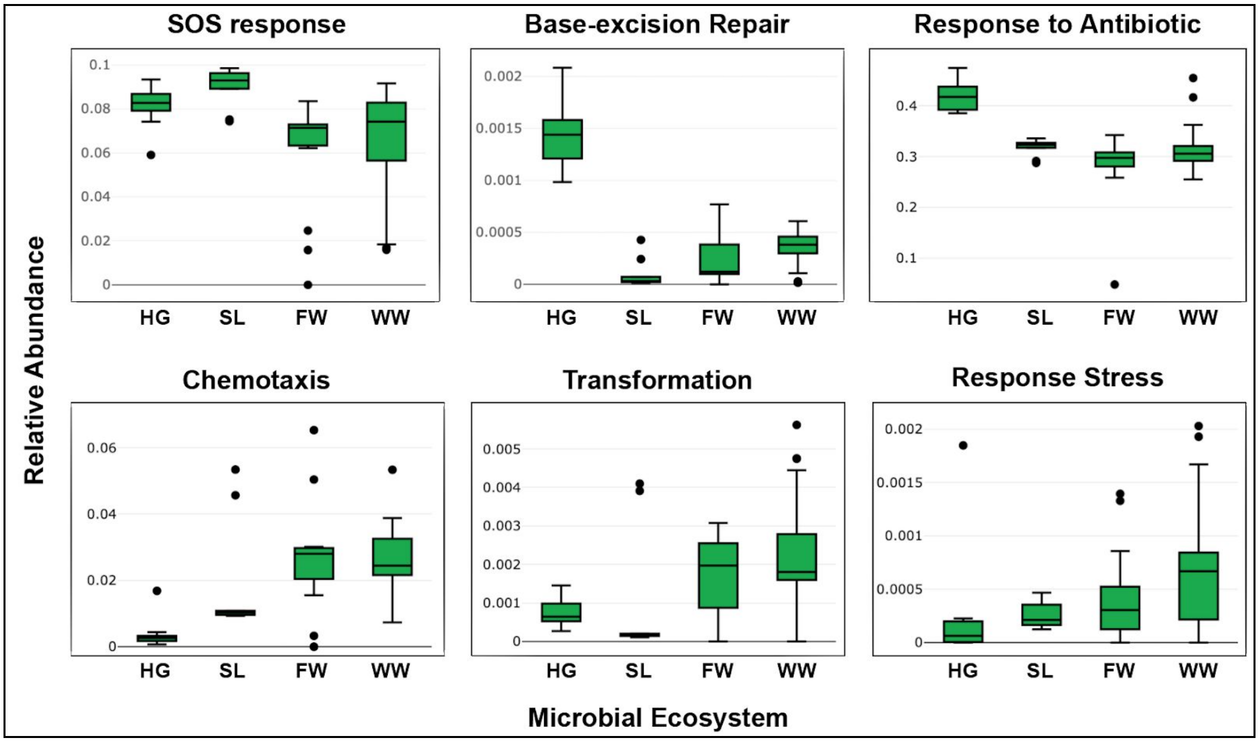

**Supplementary Figure S3**: Relative abundance of biological process from the GO response to stress database.

| Pathway | Training Proteins | Validation Genes | Validation Reads |
|---|---|---|---|
| Amino-acid_biosynthesis | 501 | 126 | 7148 |
| Amino-acid_degradation | 99 | 25 | 1422 |
| Antibiotic_biosynthesis | 82 | 20 | 1121 |
| Aromatic_compound_metabolism | 68 | 17 | 890 |
| Bacterial_outer_membrane_biogenesis | 62 | 16 | 901 |
| Carbohydrate_biosynthesis | 79 | 20 | 1071 |
| Carbohydrate_degradation | 167 | 42 | 2199 |
| Carbohydrate_metabolism | 138 | 35 | 1903 |
| Cell_wall_biogenesis | 190 | 48 | 2642 |
| Cofactor_biosynthesis | 344 | 86 | 5082 |
| Isoprenoid_biosynthesis | 42 | 10 | 651 |
| Lipid_metabolism | 177 | 45 | 2569 |

| | | | |
|---|---|---|---|
| Metabolic_intermediate_biosynthesis | 95 | 24 | 1379 |
| Nitrogen_metabolism | 50 | 12 | 709 |
| Nucleotide-sugar_biosynthesis | 42 | 10 | 505 |
| One-carbon_metabolism | 45 | 11 | 588 |
| Porphyrin-containing_compound_metabolism | 66 | 16 | 713 |
| Protein_modification | 46 | 12 | 663 |
| Purine_metabolism | 133 | 33 | 1720 |
| Pyrimidine_metabolism | 105 | 26 | 1397 |
| Xenobiotic_degradation | 41 | 10 | 478 |

**Supplementary Table S1:** UniProt pathway database with number of proteins used for training, number of genes used for validation and the simulated number of reads for each pathway category.

| **Antibiotic Class** | **Proteins** |
|---|---|
| multidrug | 4456 |
| beta-lactam | 2885 |
| MLS | 1710 |
| tetracycline | 557 |
| fosfomycin | 434 |
| aminoglycoside | 403 |
| glycopeptide | 346 |
| unclassified | 311 |
| bacitracin | 280 |
| polymyxin | 245 |
| fluoroquinolone | 158 |
| phenicol | 157 |
| sulfonamide | 125 |
| diaminopyrimidine | 80 |

**Supplementary Table S2**: Antibiotic resistance categories from ARGminer

| GO Term | Biological Process | Proteins |
|---|---|---|
| GO:0006935 | chemotaxis | 312 |
| GO:0006515 | protein_quality_control_for_misfolded_or_incompletely_synthesized_proteins | 123 |
| GO:0006298 | mismatch_repair | 1306 |
| GO:0042742 | defense_response_to_bacterium | 136 |
| GO:0046677 | response_to_antibiotic | 975 |
| GO:0009432 | SOS_response | 1030 |
| GO:0006814 | sodium_ion_transport | 125 |
| GO:0051607 | defense_response_to_virus | 149 |
| GO:0051775 | response_to_redox_state | 164 |
| GO:0045454 | cell_redox_homeostasis | 164 |
| GO:0009236 | cobalamin_biosynthetic_process | 505 |
| GO:0045910 | negative_regulation_of_DNA_recombination | 150 |
| GO:0006281 | DNA_repair | 3450 |
| GO:0006261 | DNA-dependent_DNA_replication | 228 |
| GO:0006289 | nucleotide-excision_repair | 1485 |
| GO:0010038 | response_to_metal_ion | 123 |
| GO:0009163 | nucleoside_biosynthetic_process | 204 |
| GO:0006541 | glutamine_metabolic_process | 258 |
| GO:0030091 | protein_repair | 186 |
| GO:0034605 | cellular_response_to_heat | 128 |
| GO:0019835 | cytolysis | 140 |
| GO:0006355 | regulation_of_transcription,_DNA-templated | 360 |
| GO:0019380 | 3-phenylpropionate_catabolic_process | 224 |
| GO:0045892 | negative_regulation_of_transcription,_DNA-templated | 203 |
| GO:0000724 | double-strand_break_repair_via_homologous_recombination | 296 |
| GO:0000160 | phosphorelay_signal_transduction_system | 449 |
| GO:0006979 | response_to_oxidative_stress | 677 |
| GO:0006310 | DNA_recombination | 1766 |
| GO:0043571 | maintenance_of_CRISPR_repeat_elements | 101 |
| GO:0009307 | DNA_restriction-modification_system | 283 |

| | | |
|---|---|---|
| GO:0006284 | base-excision_repair | 1097 |
| GO:0006974 | cellular_response_to_DNA_damage_stimulus | 118 |
| GO:0042744 | hydrogen_peroxide_catabolic_process | 371 |
| GO:0005975 | carbohydrate_metabolic_process | 131 |
| GO:0006260 | DNA_replication | 958 |
| GO:0009636 | response_to_toxic_substance | 149 |
| GO:0009405 | pathogenesis | 116 |
| GO:0006811 | ion_transport | 102 |
| GO:0006109 | regulation_of_carbohydrate_metabolic_process | 214 |

**Supplementary Table S3**: Database of response to stress associated categories using Gene Ontology terms.

| **Pathway** | **Precision** | **Recall** | **F1 Score** |
|---|---|---|---|
| Amino-acid_biosynthesis | 0.99 | 0.99 | 0.99 |
| Amino-acid_degradation | 0.97 | 0.97 | 0.97 |
| Antibiotic_biosynthesis | 0.93 | 0.82 | 0.87 |
| Aromatic_compound_metabolism | 0.42 | 0.44 | 0.43 |
| Bacterial_outer_membrane_biogenesis | 1 | 0.36 | 0.53 |
| Carbohydrate_biosynthesis | 0.97 | 1 | 0.98 |
| Carbohydrate_degradation | 0.99 | 0.99 | 0.99 |
| Carbohydrate_metabolism | 0.97 | 0.97 | 0.97 |
| Cell_wall_biogenesis | 0.99 | 1 | 0.99 |
| Cofactor_biosynthesis | 0.99 | 0.99 | 0.99 |
| Isoprenoid_biosynthesis | 1 | 0.99 | 0.99 |
| Lipid_metabolism | 0.98 | 1 | 0.99 |
| Metabolic_intermediate_biosynthesis | 0.97 | 0.97 | 0.97 |
| Nitrogen_metabolism | 1 | 1 | 1 |
| Nucleotide-sugar_biosynthesis | 1 | 0.99 | 1 |
| One-carbon_metabolism | 0.99 | 0.95 | 0.97 |
| Porphyrin-containing_compound_metabolism | 0.99 | 0.97 | 0.98 |
| Protein_modification | 1 | 1 | 1 |
| Purine_metabolism | 1 | 0.99 | 0.99 |

| | | | |
|---|---|---|---|
| Pyrimidine_metabolism | 0.99 | 1 | 1 |
| Xenobiotic_degradation | 0.8 | 0.46 | 0.59 |
| **Average** | **0.99** | **0.99** | **0.99** |

**Supplementary Table S5**: Prediction performance of MetaMLP.

| **Pathway** | **Precision** | **Recall** | **F1 Score** |
|---|---|---|---|
| Amino-acid_biosynthesis | 1 | 1 | 1 |
| Amino-acid_degradation | 0.94 | 1 | 0.97 |
| Antibiotic_biosynthesis | 1 | 0.97 | 0.98 |
| Aromatic_compound_metabolism | 0.78 | 1 | 0.87 |
| Bacterial_outer_membrane_biogenesis | 0 | 0 | 0 |
| Carbohydrate_biosynthesis | 1 | 0.98 | 0.99 |
| Carbohydrate_degradation | 1 | 1 | 1 |
| Carbohydrate_metabolism | 0.97 | 1 | 0.99 |
| Cell_wall_biogenesis | 1 | 1 | 1 |
| Cofactor_biosynthesis | 1 | 1 | 1 |
| Isoprenoid_biosynthesis | 1 | 1 | 1 |
| Lipid_metabolism | 1 | 1 | 1 |
| Metabolic_intermediate_biosynthesis | 1 | 1 | 1 |
| Nitrogen_metabolism | 1 | 1 | 1 |
| Nucleotide-sugar_biosynthesis | 1 | 1 | 1 |
| One-carbon_metabolism | 1 | 1 | 1 |
| Porphyrin-containing_compound_metabolism | 1 | 1 | 1 |
| Protein_modification | 1 | 1 | 1 |
| Purine_metabolism | 1 | 1 | 1 |
| Pyrimidine_metabolism | 1 | 1 | 1 |
| Xenobiotic_degradation | 1 | 0.47 | 0.64 |
| **Average** | 0.99 | 1 | 1 |

**Supplementary Table S6**: Prediction performance of the best hit approach using diamond blastx.
