## Supplementary table S4 for "MetaMLP: A fast word embedding based classifier to profile target gene databases in metagenomic samples"

| Type | Experiment Accession | Instrument | Submitter | Study Accession | Study Title | Sample Accession | Total Size, Mb | Total Spots | Total Bases | Library Strategy | Library Source |
|---|---|---|---|---|---|---|---|---|---|---|---|
| wastewater | SRX425466 | Illumina HiSeq 20 | Nanjing University | SRP035334 | Illumina shotgun sequencing reads from four municipal sewage wastewater trea | SRS529890 | 599.26 | 12147592 | 1226906792 | WGS | METAGENOMIC |
| wastewater | SRX425457 | Illumina HiSeq 20 | Nanjing University | SRP035334 | Illumina shotgun sequencing reads from four municipal sewage wastewater trea | SRS529882 | 620.79 | 12472820 | 1259754820 | WGS | METAGENOMIC |
| freshwater | SRX4384578 | Illumina HiSeq 25 | National Research Council Canada | SRP151914 | NRC/UOttawa - Peace Athabasca Delta Raw sequence reads | SRS3540779 | 838.11 | 10068187 | 2517046750 | WGS | METAGENOMIC |
| freshwater | SRX4384610 | Illumina HiSeq 25 | National Research Council Canada | SRP151914 | NRC/UOttawa - Peace Athabasca Delta Raw sequence reads | SRS3540807 | 860.23 | 10071510 | 2517877500 | WGS | METAGENOMIC |
| wastewater | SRX2830179 | Illumina HiSeq 25 | Northwestern University | SRP107015 | Wastewater, sludge and lake sediment metagenomes Raw sequence reads | SRS2205682 | 939.12 | 14610430 | 2922086000 | WGS | METAGENOMIC |
| freshwater | SRX4384590 | Illumina HiSeq 25 | National Research Council Canada | SRP151914 | NRC/UOttawa - Peace Athabasca Delta Raw sequence reads | SRS3540828 | 967.33 | 11031126 | 2757781500 | WGS | METAGENOMIC |
| wastewater | SRX2830180 | Illumina HiSeq 25 | Northwestern University | SRP107015 | Wastewater, sludge and lake sediment metagenomes Raw sequence reads | SRS2205683 | 985.5 | 15282996 | 3056599200 | WGS | METAGENOMIC |
| wastewater | SRX2830177 | Illumina HiSeq 25 | Northwestern University | SRP107015 | Wastewater, sludge and lake sediment metagenomes Raw sequence reads | SRS2205680 | 1004.43 | 15559978 | 3111995600 | WGS | METAGENOMIC |
| freshwater | SRX4384596 | Illumina HiSeq 25 | National Research Council Canada | SRP151914 | NRC/UOttawa - Peace Athabasca Delta Raw sequence reads | SRS3540795 | 1084.01 | 12600830 | 3150207500 | WGS | METAGENOMIC |
| freshwater | SRX4384588 | Illumina HiSeq 25 | National Research Council Canada | SRP151914 | NRC/UOttawa - Peace Athabasca Delta Raw sequence reads | SRS3540789 | 1098.05 | 12549678 | 3137419500 | WGS | METAGENOMIC |
| freshwater | SRX4384595 | Illumina HiSeq 25 | National Research Council Canada | SRP151914 | NRC/UOttawa - Peace Athabasca Delta Raw sequence reads | SRS3540794 | 1198.54 | 13817712 | 3454428000 | WGS | METAGENOMIC |
| Human | SRX5023882 | Illumina HiSeq 40 | Oregon State University | SRP169523 | human gut metagenome Metagenome | SRS4055733 | 1310.88 | 10045503 | 3033741906 | WGS | METAGENOMIC |
| Human | SRX5023884 | Illumina HiSeq 40 | Oregon State University | SRP169523 | human gut metagenome Metagenome | SRS4055735 | 1366.17 | 10721856 | 3238000512 | WGS | METAGENOMIC |
| freshwater | SRX4384609 | Illumina HiSeq 25 | National Research Council Canada | SRP151914 | NRC/UOttawa - Peace Athabasca Delta Raw sequence reads | SRS3540806 | 1371.16 | 15417471 | 3854367750 | WGS | METAGENOMIC |
| freshwater | SRX4384584 | Illumina HiSeq 25 | National Research Council Canada | SRP151914 | NRC/UOttawa - Peace Athabasca Delta Raw sequence reads | SRS3540784 | 1373.9 | 16475111 | 4118777750 | WGS | METAGENOMIC |
| freshwater | SRX4384579 | Illumina HiSeq 25 | National Research Council Canada | SRP151914 | NRC/UOttawa - Peace Athabasca Delta Raw sequence reads | SRS3540781 | 1376.29 | 15682558 | 3920639500 | WGS | METAGENOMIC |
| Human | SRX5023912 | Illumina HiSeq 40 | Oregon State University | SRP169523 | human gut metagenome Metagenome | SRS4055763 | 1394.88 | 10125103 | 3057781106 | WGS | METAGENOMIC |
| Human | SRX5023883 | Illumina HiSeq 40 | Oregon State University | SRP169523 | human gut metagenome Metagenome | SRS4055734 | 1417.31 | 10875631 | 3284440562 | WGS | METAGENOMIC |
| freshwater | SRX4384606 | Illumina HiSeq 25 | National Research Council Canada | SRP151914 | NRC/UOttawa - Peace Athabasca Delta Raw sequence reads | SRS3540804 | 1424.02 | 16160830 | 4040207500 | WGS | METAGENOMIC |
| Human | SRX5023885 | Illumina HiSeq 40 | Oregon State University | SRP169523 | human gut metagenome Metagenome | SRS4055736 | 1462.96 | 11132135 | 3361904770 | WGS | METAGENOMIC |
| Human | SRX5023877 | Illumina HiSeq 40 | Oregon State University | SRP169523 | human gut metagenome Metagenome | SRS4055728 | 1480.19 | 10767883 | 3251900666 | WGS | METAGENOMIC |
| Human | SRX5023907 | Illumina HiSeq 40 | Oregon State University | SRP169523 | human gut metagenome Metagenome | SRS4055758 | 1552.07 | 11213465 | 3386466430 | WGS | METAGENOMIC |
| Human | SRX5023913 | Illumina HiSeq 40 | Oregon State University | SRP169523 | human gut metagenome Metagenome | SRS4055764 | 1617.26 | 12479814 | 3768903828 | WGS | METAGENOMIC |
| wastewater | ERX1054406 | Illumina HiSeq 20 | AALBORG UNIVERSITY | ERP011345 | Retrieval of Commamox genomes using metagenomics | ERS805489 | 1634.84 | 13956081 | 2791216200 | WGS | METAGENOMIC |
| Human | SRX5023902 | Illumina HiSeq 40 | Oregon State University | SRP169523 | human gut metagenome Metagenome | SRS4055753 | 1770.16 | 13839737 | 4179600574 | WGS | METAGENOMIC |
| Human | SRX5023910 | Illumina HiSeq 40 | Oregon State University | SRP169523 | human gut metagenome Metagenome | SRS4055761 | 1782.97 | 13710260 | 4140498520 | WGS | METAGENOMIC |
| Human | SRX5023890 | Illumina HiSeq 40 | Oregon State University | SRP169523 | human gut metagenome Metagenome | SRS4055741 | 1888.02 | 14269837 | 4309490774 | WGS | METAGENOMIC |
| Human | SRX5023879 | Illumina HiSeq 40 | Oregon State University | SRP169523 | human gut metagenome Metagenome | SRS4055730 | 2236.78 | 16777266 | 5066734332 | WGS | METAGENOMIC |
| wastewater | ERX656380 | Illumina HiSeq 20 | AALBORG UNIVERSITY | ERP009124 | Metagenomes of Danish EBPR WWTPs | ERS632917 | 2352.65 | 13173900 | 3952170000 | WGS | METAGENOMIC |
| Human | SRX5023904 | Illumina HiSeq 40 | Oregon State University | SRP169523 | human gut metagenome Metagenome | SRS4055755 | 2379.03 | 18032370 | 5445775740 | WGS | METAGENOMIC |
| Human | SRX5023900 | Illumina HiSeq 40 | Oregon State University | SRP169523 | human gut metagenome Metagenome | SRS4055751 | 2518.18 | 18213008 | 5500328416 | WGS | METAGENOMIC |
| wastewater | SRX3189259 | Illumina HiSeq 15 | Karlsruhe Institute of Technology | SRP117738 | Live-Dead discrimination analysis is essential for molecular biology evaluation o | SRS2516594 | 2669.3 | 31365642 | 6273128400 | WGS | METAGENOMIC |
| Human | SRX5023906 | Illumina HiSeq 40 | Oregon State University | SRP169523 | human gut metagenome Metagenome | SRS4055757 | 2731.6 | 19886441 | 6005705182 | WGS | METAGENOMIC |
| wastewater | SRX3189261 | Illumina HiSeq 15 | Karlsruhe Institute of Technology | SRP117738 | Live-Dead discrimination analysis is essential for molecular biology evaluation o | SRS2516596 | 2739.27 | 32192124 | 6438424800 | WGS | METAGENOMIC |
| Soil | SRX1272842 | Illumina HiSeq 25 | Tsinghua University | SRP063891 | HP and HPP metagenome comparison | SRS1079806 | 2904.55 | 20967536 | 5241884000 | WGS | METAGENOMIC |
| Soil | SRX1264433 | Illumina HiSeq 25 | Tsinghua University | SRP063891 | HP and HPP metagenome comparison | SRS1074972 | 2981.15 | 22590236 | 5647559000 | WGS | METAGENOMIC |
| wastewater | ERX1054407 | Illumina HiSeq 20 | AALBORG UNIVERSITY | ERP011345 | Retrieval of Commamox genomes using metagenomics | ERS805490 | 3054.72 | 26057980 | 5211596000 | WGS | METAGENOMIC |
| freshwater | SRX4015216 | Illumina HiSeq 25 | andrew_boddicker's shared submissic | SRP144089 | freshwater metagenome Genome sequencing and assembly | SRS3236713 | 3226.61 | 33376046 | 8410763592 | WGS | METAGENOMIC |
| wastewater | SRX3189260 | Illumina HiSeq 15 | Karlsruhe Institute of Technology | SRP117738 | Live-Dead discrimination analysis is essential for molecular biology evaluation o | SRS2516595 | 3318.9 | 39340492 | 7868098400 | WGS | METAGENOMIC |
| wastewater | SRX451135 | Illumina HiSeq 25 | Tongji Univ | SRP035848 | Metagenomic datasets of activated sludge samples | SRS544436 | 3530.02 | 29167563 | 5891847726 | WGS | METAGENOMIC |
| wastewater | SRX451093 | Illumina HiSeq 25 | Tongji Univ | SRP035848 | Metagenomic datasets of activated sludge samples | SRS544428 | 3711.98 | 30702043 | 6201812686 | WGS | METAGENOMIC |
| freshwater | SRX4015219 | Illumina HiSeq 25 | andrew_boddicker's shared submissic | SRP144089 | freshwater metagenome Genome sequencing and assembly | SRS3236718 | 4173.58 | 43549092 | 10974371184 | WGS | METAGENOMIC |
| wastewater | SRX451165 | Illumina HiSeq 25 | Tongji Univ | SRP035848 | Metagenomic datasets of activated sludge samples | SRS544439 | 4454.93 | 36873995 | 7448546990 | WGS | METAGENOMIC |
| wastewater | SRX450092 | Illumina HiSeq 25 | Tongji Univ | SRP035848 | Metagenomic datasets of activated sludge samples | SRS543595 | 4457.88 | 35438248 | 7141549089 | WGS | METAGENOMIC |
| wastewater | ERX1054408 | Illumina HiSeq 20 | AALBORG UNIVERSITY | ERP011345 | Retrieval of Commamox genomes using metagenomics | ERS805491 | 4785.09 | 41133102 | 8226620400 | WGS | METAGENOMIC |
| wastewater | ERX656401 | Illumina HiSeq 20 | AALBORG UNIVERSITY | ERP009124 | Metagenomes of Danish EBPR WWTPs | ERS632931 | 8132.37 | 42999422 | 12899826600 | WGS | METAGENOMIC |
| freshwater | SRX4478628 | Illumina HiSeq 25 | JGI | SRP155578 | Freshwater microbial communities from McNutts Creek, Athens, Georgia, Unitec | SRS3603646 | 8626.29 | 59373376 | 17930759552 | WGS | METAGENOMIC |
| Soil | SRX1299772 | Illumina HiSeq 20 | University of California, Berkeley | SRP064390 | Meadow Soil samples from Angelo, CA Genome sequencing and assembly | SRS1096859 | 9003.87 | 31492291 | 15809130082 | WGS | METAGENOMIC |
| Soil | SRX1299765 | Illumina HiSeq 20 | University of California, Berkeley | SRP064390 | Meadow Soil samples from Angelo, CA Genome sequencing and assembly | SRS1096866 | 9977.6 | 35885821 | 18014682142 | WGS | METAGENOMIC |
| wastewater | ERX656379 | Illumina HiSeq 20 | AALBORG UNIVERSITY | ERP009124 | Metagenomes of Danish EBPR WWTPs | ERS632916 | 10094.27 | 59871310 | 17961393000 | WGS | METAGENOMIC |
| wastewater | ERX656391 | Illumina HiSeq 20 | AALBORG UNIVERSITY | ERP009124 | Metagenomes of Danish EBPR WWTPs | ERS632922 | 10878.54 | 54568514 | 16370554200 | WGS | METAGENOMIC |
| Soil | SRX1299769 | Illumina HiSeq 20 | University of California, Berkeley | SRP064390 | Meadow Soil samples from Angelo, CA Genome sequencing and assembly | SRS1096862 | 11186.85 | 40858831 | 20511133162 | WGS | METAGENOMIC |
| Soil | SRX1299770 | Illumina HiSeq 20 | University of California, Berkeley | SRP064390 | Meadow Soil samples from Angelo, CA Genome sequencing and assembly | SRS1096860 | 11406.61 | 40484382 | 20323159764 | WGS | METAGENOMIC |
| Soil | SRX1299768 | Illumina HiSeq 20 | University of California, Berkeley | SRP064390 | Meadow Soil samples from Angelo, CA Genome sequencing and assembly | SRS1096863 | 11745.36 | 42670900 | 21420791800 | WGS | METAGENOMIC |
| freshwater | SRX4807264 | Illumina HiSeq 25 | JGI | SRP164129 | Freshwater microbial communities from Pennsylvania, USA, analyzing microbe ( | SRS3885806 | 11784.97 | 77276991 | 23337651282 | WGS | METAGENOMIC |
| wastewater | ERX656392 | Illumina HiSeq 20 | AALBORG UNIVERSITY | ERP009124 | Metagenomes of Danish EBPR WWTPs | ERS632923 | 11931.4 | 60065658 | 18019697400 | WGS | METAGENOMIC |
| freshwater | SRX4807070 | Illumina HiSeq 25 | JGI | SRP163939 | Freshwater microbial communities from Pennsylvania, USA, analyzing microbe ( | SRS3885617 | 11947.7 | 80865366 | 24421340532 | WGS | METAGENOMIC |
| Soil | SRX1299764 | Illumina HiSeq 20 | University of California, Berkeley | SRP064390 | Meadow Soil samples from Angelo, CA Genome sequencing and assembly | SRS1096867 | 12018.83 | 43399517 | 21786557534 | WGS | METAGENOMIC |
| Soil | SRX1299771 | Illumina HiSeq 20 | University of California, Berkeley | SRP064390 | Meadow Soil samples from Angelo, CA Genome sequencing and assembly | SRS1096861 | 12184.68 | 40983076 | 20573504152 | WGS | METAGENOMIC |
| wastewater | ERX656395 | Illumina HiSeq 20 | AALBORG UNIVERSITY | ERP009124 | Metagenomes of Danish EBPR WWTPs | ERS632925 | 12358.81 | 67619077 | 20285723100 | WGS | METAGENOMIC |
| wastewater | ERX656398 | Illumina HiSeq 20 | AALBORG UNIVERSITY | ERP009124 | Metagenomes of Danish EBPR WWTPs | ERS632928 | 12462.12 | 64887815 | 19466344500 | WGS | METAGENOMIC |
| wastewater | ERX656390 | Illumina HiSeq 20 | AALBORG UNIVERSITY | ERP009124 | Metagenomes of Danish EBPR WWTPs | ERS632921 | 12673.88 | 72877460 | 21863238000 | WGS | METAGENOMIC |
| wastewater | ERX656400 | Illumina HiSeq 20 | AALBORG UNIVERSITY | ERP009124 | Metagenomes of Danish EBPR WWTPs | ERS632930 | 15049.52 | 82409629 | 24722888700 | WGS | METAGENOMIC |
| Soil | SRX1299773 | Illumina HiSeq 20 | University of California, Berkeley | SRP064390 | Meadow Soil samples from Angelo, CA Genome sequencing and assembly | SRS1096858 | 16212.05 | 57695413 | 28963097326 | WGS | METAGENOMIC |

| | | | | | | | | | | | |
|---|---|---|---|---|---|---|---|---|---|---|---|
| wastewater | ERX656378 | Illumina HiSeq 20 | AALBORG UNIVERSITY | ERP009124 | Metagenomes of Danish EBPR WWTPs | ERS632915 | 16305.99 | 89347369 | 26804210700 | WGS | METAGENOMIC |
| wastewater | ERX656397 | Illumina HiSeq 20 | AALBORG UNIVERSITY | ERP009124 | Metagenomes of Danish EBPR WWTPs | ERS632927 | 16528.92 | 86815483 | 26044644900 | WGS | METAGENOMIC |
| wastewater | ERX656396 | Illumina HiSeq 20 | AALBORG UNIVERSITY | ERP009124 | Metagenomes of Danish EBPR WWTPs | ERS632926 | 19568.24 | 98562152 | 29568645600 | WGS | METAGENOMIC |
